# Identification of significant molecules and signaling pathways between 2D and 3D culture methods of renal cancer cells

**DOI:** 10.1101/2022.01.31.478529

**Authors:** Mengyao Wang, Hongmei Guo, Hanming Gu, Mason Zhang

**Author notes:** Corresponding author: Hanming Gu, SHU-UTS SILC School, Shanghai University, Shanghai, China.

## Abstract

The majority of cancer studies are conducted with the two-dimensional (2D) culture method, which does not reflect tumor in vivo structure. The 3D culture method can form free bundles of cancer cells and spheroid, which mimics the tumor microenvironment in vivo. However, the molecules and signaling pathways between the 2D and 3D culture methods are still unknown. In this study, we aim to identify the key molecules and signaling pathways by analyzing the RNA-seq data. The GSE190296 was created by the BGISEQ-500 (Homo sapiens). The KEGG and GO analyses indicated sulfur compound metabolic process and regulation of leukocyte mediated immunity are the major differences between 2D and 3D renal cancer cell cultures. Moreover, we figured out several interactive genes including MYC, EGF, VEGFA, STAT3, NOTCH1, CAT, CCND1, HSPA8, DLG4, and HSPA5. Our study may provide new knowledge on the differences between 2D and 3D cancer cell cultures.

## Introduction

In the US and Europe, renal cancer was ranked as the high common malignancy in men each year^1^. Though surgery is an effective therapy for early diagnosed patients, a number of patients suffer recurrence or metastatic cancers thereafter^2^. Limited treatment methods have been doable for these patients. The classic signs of pain, hematuria and flank mass are indicative of advanced diseases. There are about 30% of patients with renal cancer showed metastatic diseases, and 45% with localized diseases^3^.

Radical nephrectomy is the standard method for renal cancer and no chemotherapeutic regimen is considered as a standard method^4^. Improvements in molecular knowledge of resistance have contributed to the identification of new drug targets^5^. Many candidate agents are under evaluation, which may play a vital role in combination therapy^6^. One of the important factors in vitro studies is the culture method^7^. Traditionally, the anti-cancer drugs have been tested in 2D cultured cancer cells, but this culture method is not suitable for determining the tumor microenvironment created by the 3D culture method^8^. Thus, the 3D culture method is better than the 2D culture method to mimic the true environment of renal cancer.

In this study, we compared and analyzed the methods of cancer cells culture by using the RNA-seq data. We discovered a number of DEGs and significant signaling pathways. We then performed the gene enrichment and protein-protein interaction (PPI) network analysis to obtain the interacting map and key genes. These genes and biological processes may provide new knowledge of renal cancer cells in different environments.

## Methods

### Data resources

Gene dataset GSE190296 was downloaded from the GEO database. The data was produced by the Illumina NextSeq 500 (Homo sapiens) (Military Institute of Medicine, Szaserow 128, Warsaw, Poland). The analyzed dataset includes three RenCa cells cultured by the 2D method and three RenCa cells cultured by the 3D method.

### Data acquisition and processing

The data were organized and conducted by the R package as previously described^9–13^. We used a classical t-test to identify DEGs with P<0.05 and fold change ≥1.5 as being statistically significant.

The Kyoto Encyclopedia of Genes and Genomes (KEGG) and Gene Ontology (GO) KEGG and GO analyses were performed by the R package (ClusterProfiler) and Reactome^12, 14, 15^. P<0.05 was considered statistically significant.

### Protein-protein interaction (PPI) networks

The Molecular Complex Detection (MCODE) was used to create the PPI networks. The significant modules were produced from constructed PPI networks and String networks (https://string-db.org/). The biological processes analyses were performed by using Reactome (https://reactome.org/), and P<0.05 was considered significant.

## Results

### Identification of DEGs in renal cancer cells between 2D and 3D culture systems

To determine the effects of different cancer cell culture systems, we analyzed the RNA-seq data of renal cancer cells from 2D and 3D culture systems. A total of 1281 genes were identified with the threshold of P < 0.01. The top up- and down-regulated genes were shown by the heatmap and volcano plot (Figure 1). The top ten DEGs were listed in Table 1.

**Figure 1.**
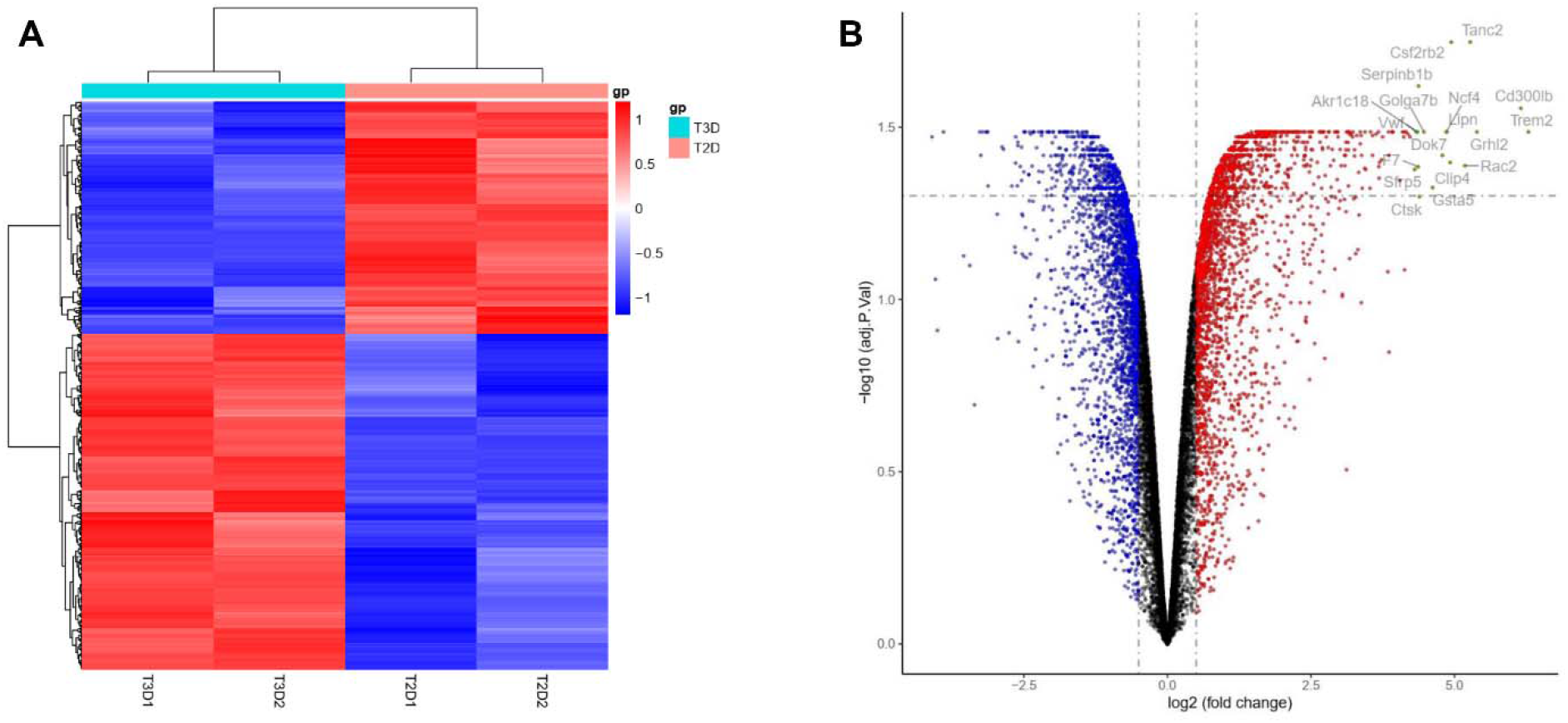
Heatmap and volcano plot in renal cancer cells between 2D and 3D culture systems. (A) Heatmap of significant DEGs. Significant DEGs (P < 0.05) were used to construct the heatmap. T2D, tumor cells of 2D culture; T3D, tumor cells of 3D culture. (B) Volcano plot for DEGs in renal cancer cells between 2D and 3D culture systems. The most significantly changed genes are highlighted by grey dots.

**Table 1.**

| **Entrez gene** | **Gene symbol** | **Fold-change** | **Regulation** |
| --- | --- | --- | --- |
| Top 10 down-regulated DEGs |  |  |  |
| 342527 | Smtnl2 | -4.098803795 | Down |
| 545363 | Gm5833 | -3.899557414 | Down |
| 2152 | F3 | -3.267920658 | Down |
| 10841 | Ftcd | -3.251459755 | Down |
| 9293 | Gpr52 | -3.219829955 | Down |
| 10252 | Spry1 | -3.157431352 | Down |
| 196410 | Mettl7b | -3.14920572 | Down |
| 85477 | Scin | -3.128462553 | Down |
| 100040843 | Cyp4a32 | -3.123319545 | Down |
| 98736 | 1700034H15Rik | -2.87893792 | Down |
| Top 10 up-regulated DEGs |  |  |  |
| 54209 | Trem2 | 6.286834421 | up |
| 124599 | Cd300lb | 6.154935072 | up |
| 79977 | Grhl2 | 5.388158156 | up |
| 26115 | Tanc2 | 5.272378718 | up |
| 5880 | Rac2 | 5.181825342 | up |
| 12984 | Csf2rb2 | 4.94316349 | up |
| 79745 | Clip4 | 4.923979988 | up |
| 643418 | Lipn | 4.865791822 | up |
| 4689 | Ncf4 | 4.85610423 | up |
| 285489 | Dok7 | 4.788819374 | up |

### Enrichment analysis of DEGs in renal cancer cells between 2D and 3D culture systems

To further understand the impacts on different culture systems in renal cancer cells, we performed the KEGG and GO enrichment (Figure 2). We figured out the top ten KEGG items including “Human cytomegalovirus infection”, “Epstein-Barr virus infection”, “Cellular senescence”, “Phagosome”, “Lysosome”, “Antigen processing and presentation”, “Glutathione metabolism”, “Platinum drug resistance”, “Viral myocarditis”, and “Drug metabolism - cytochrome P450”. We identified the top ten BP of GO items, which contains “sulfur compound metabolic process”, “Regulation of leukocyte mediated immunity”, “Cellular modified amino acid metabolic process”, “Regulation of adaptive”, “Immune response based on somatic recombination of immune receptors built from immunoglobulin superfamily domains”, “Glutathione metabolic process”, “T cell mediated cytotoxicity”, “Regulation of T cell mediated cytotoxicity”, “Antigen processing and presentation of endogenous peptide antigen”, “Antigen processing and presentation of endogenous peptide antigen via MHC class I”, and “Antigen processing and presentation of endogenous antigen”. We identified the top ten CC of GO items, which contains “Apical part of cell”, “Vacuolar membrane”, “Collagen-containing extracellular matrix”, “Apical plasma membrane”, “Early endosome”, “Endosome membrane”, “Lysosomal membrane”, “Lytic vacuole membrane”, “MHC class I protein complex”, and “Phagocytic vesicle membrane”. We also identified the top ten MF of GO items including “Amide binding”, “Peptide binding”, “Sulfur compound binding”, “Transferase activity, transferring alkyl or aryl (other than methyl) groups”, “Glutathione transferase activity”, “TAP binding”, “TAP1 binding”, “Glutathione binding”, “Oligopeptide binding”, and “TAP2 binding”.

**Figure 2.**
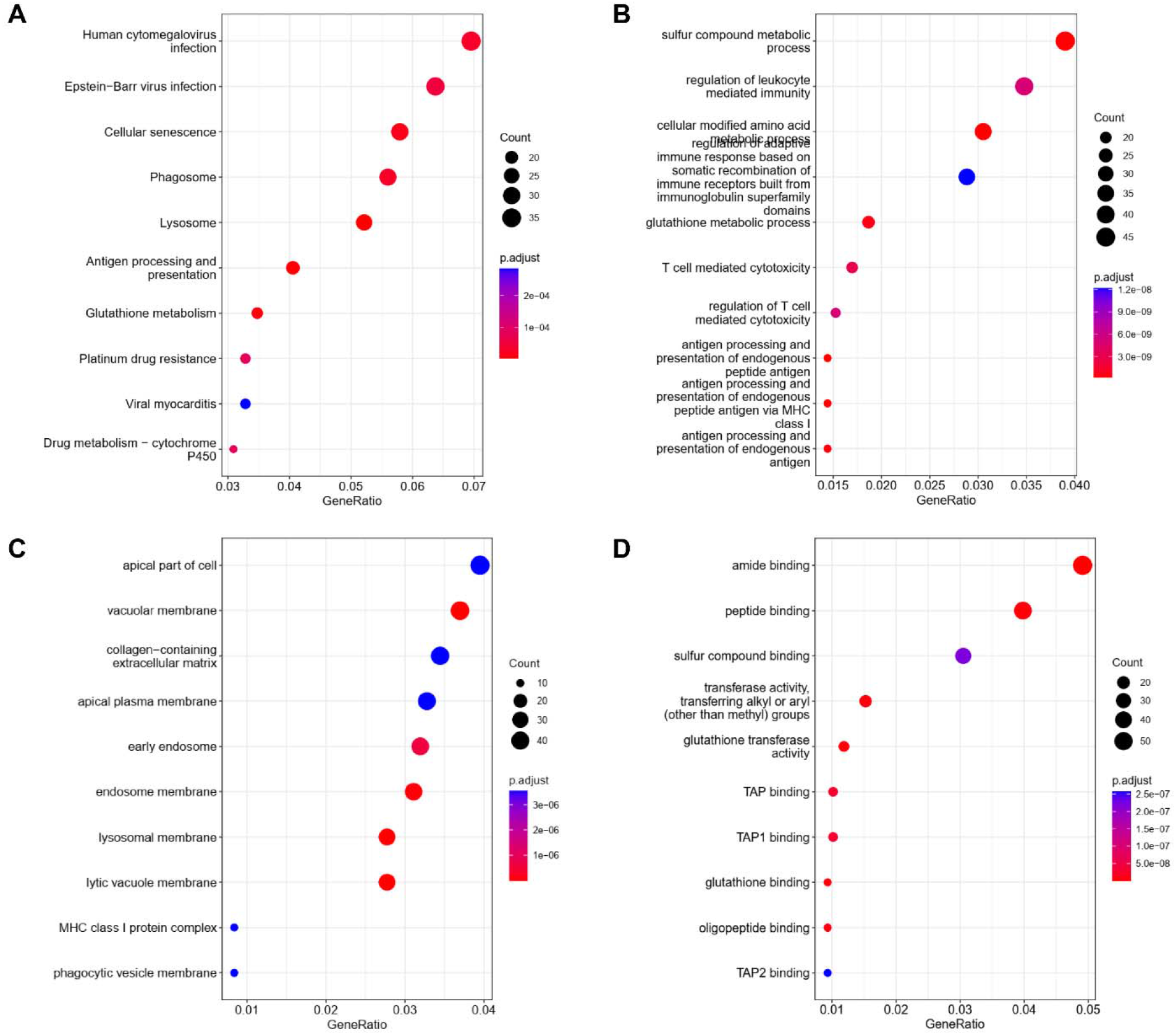
KEGG and GO analyses of DEGs in renal cancer cells between 2D and 3D culture systems. (A) KEGG analysis (B) BP: Biological processes, (C) CC: Cellular components, (D) MF: Molecular functions.

### PPI network and Reactome analyses

To explore the potential relationship among the DEGs, we created the PPI network by using 1059 nodes and 3783 edges. The combined score > 0.2 was set as a cutoff by using the Cytoscope software. Table 2 indicated the top ten genes with the highest scores. The top two significant modules were presented in Figure 3. We further analyzed the PPI and DEGs with Reactome map (Figure 4) and identified the top ten biological processes including “Glutathione conjugation”, “Response of EIF2AK1 (HRI) to heme deficiency”, “RUNX3 regulates WNT signaling”, “NGF-stimulated transcription”, “NrCAM interactions”, “Removal of aminoterminal propeptides from gamma-carboxylated proteins”, “CHL1 interactions”, “RUNX3 regulates NOTCH signaling”, “Gamma-carboxylation, transport, and amino-terminal cleavage of proteins”, and “ATF6 (ATF6-alpha) activates chaperones” (Supplemental Table S1).

**Figure 3.**
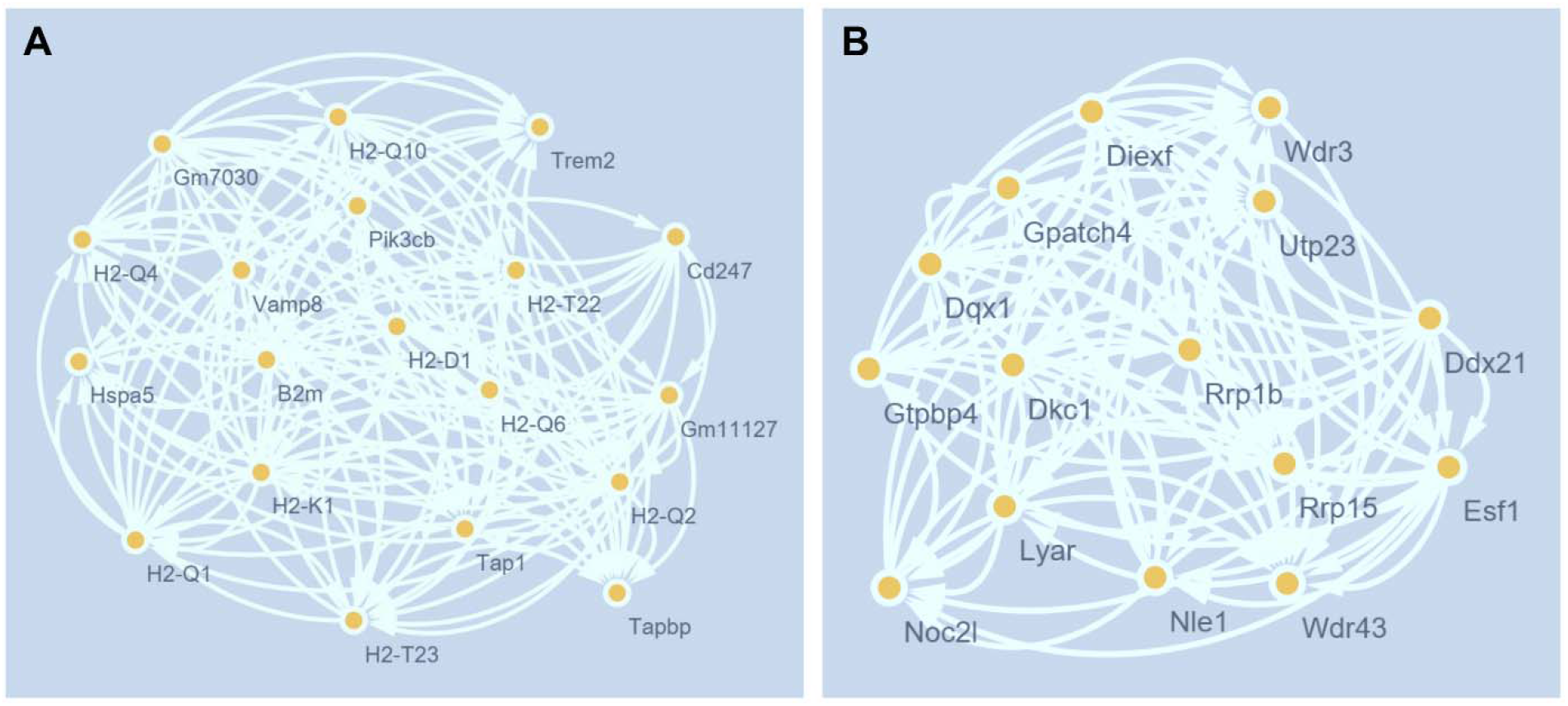
The PPI network analyses of DEGs in renal cancer cells between 2D and 3D culture systems. The cluster (A) and cluster (B) were constructed by MCODE.

**Figure 4.**
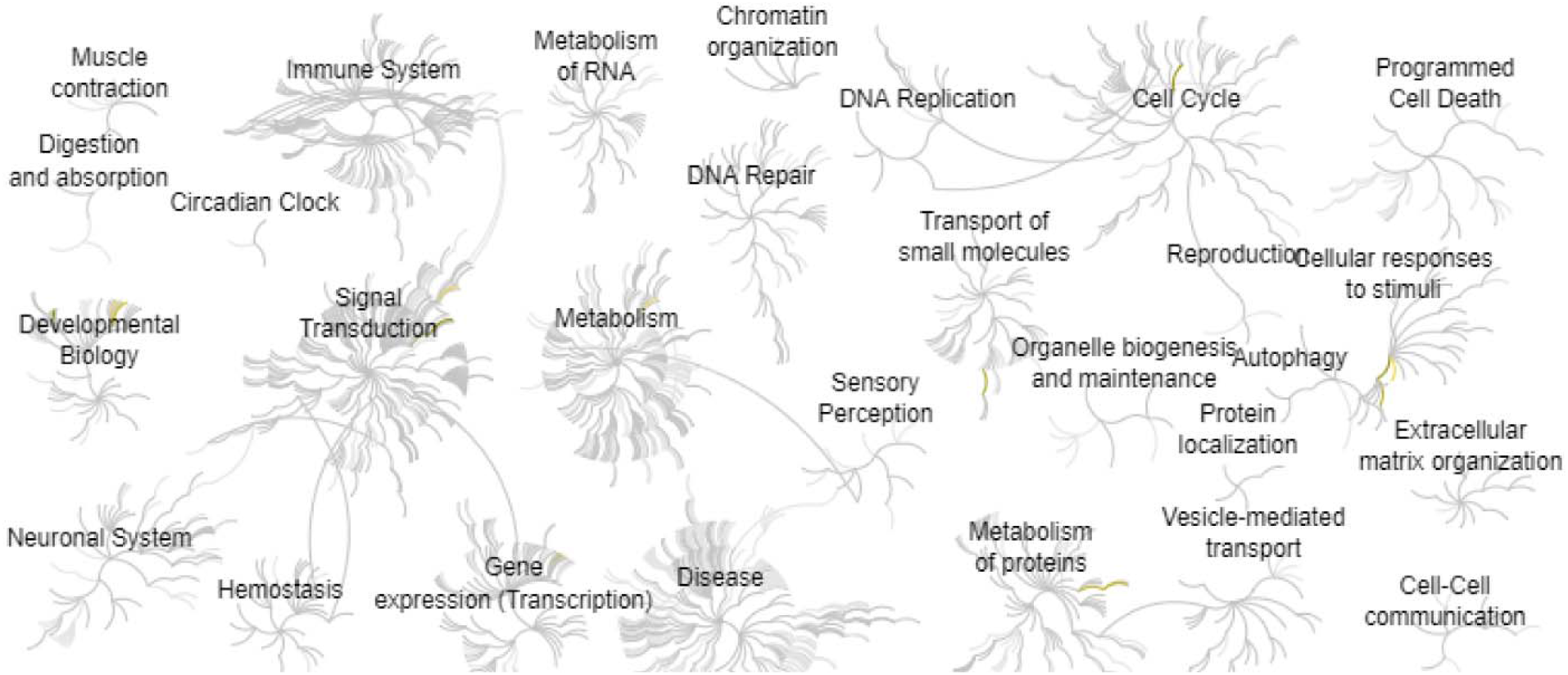
Reactome map representation of the significant biological processes in renal cancer cells between 2D and 3D culture systems.

**Table 2.** Top ten genes demonstrated by connectivity degree in the PPI network.

| **Gene symbol** | **Gene title** | **Degree** |
| --- | --- | --- |
| MYC | MYC proto-oncogene, bhlh transcription factor | 97 |
| EGF | Epidermal growth factor | 83 |
| VEGFA | Vascular endothelial growth factor A | 78 |
| STAT3 | Signal transducer and activator of transcription 3 | 68 |
| NOTCH1 | Notch receptor 1 | 59 |
| CAT | Catalase | 54 |
| CCND1 | Cyclin D1 | 52 |
| HSPA8 | Heat shock protein family A (Hsp70) member 8 | 52 |
| DLG4 | Discs large MAGUK scaffold protein 4 | 51 |
| HSPA5 | Heat shock protein family A (Hsp70) member 5 | 51 |

## Discussion

Cell culture is important for cancer research. Recently, most cells are cultured by the 2D method, but several studies showed 3D cell culture had shown improvements such as cell morphology and number monitoring, proliferation, differentiation, and drug metabolism^16^. However, the molecular difference of 2D and 3D culture is still not clear.

By analyzing the KEGG and GO enrichment, we found that “sulfur compound metabolic process” and “regulation of leukocyte mediated immunity” are the major differences between 2D and 3D renal cancer cell culture. Sulfur metabolism is controlled by LIAS that is required for the functions of the tricarboxylic acid (TCA) cycle, which stimulates mitochondrial oxidative metabolism and cancer progression. The disorder of metabolism is a potential hallmark of cancer. Nathan P Ward et al found that sulfur metabolism is controlled by LIAS that is required for the functions of the tricarboxylic acid (TCA) cycle, which stimulates mitochondrial oxidative metabolism and cancer progression^17, 18^. Moreover, Mariarita Brancaccio et al found that sulfur compounds can inhibit the activity of γ-glutamyl transpeptidase in cancer cells^19^. Renal cancer is considered an immunogenic tumor, which involves the mediate immune dysfunction by eliciting the immune cells including T cells and macrophages into the tumor microenvironment^20^. Andrea Muto et al reported the novel renal cancer therapy by targeting the immune checkpoint inhibitors^21^.

In this study, we also identified several interactive molecules by comparing the 2D and 3D methods in culturing renal cancer cells. Emelyn H Shroff et al found the overexpression of MYC promotes renal cancers through glutamine metabolism^22^. Claire Bouvard et al also found inhibition of MYC by small molecules can inhibit cancer growth in rodent xenograft models^23^. The circadian clocks and their downstream factors are responsible for a variety of physiological and pathological functions including metabolism, aging, and immunity^24–34^. Interestingly, Jamison B Burchett et al found that MYC in cancer cells can disturb the molecular clock, whereas the clock disruption in cancer can promote MYC^35^. Justin P Favaro et al found that the epidermal growth factor (EGF) is critical for tumor progression in renal cancers, which can be considered as a potential drug target^36^. VEGFA is an important mediator of angiogenesis and plays a vital role in cancer angiogenesis and progression^37^. STAT3 has a crucial role in the progression of different cancers and its activation is identified as an important target for cancer therapy^38^. GPCR/RGS signaling regulates the majority of biological processes including immunity, metabolism, aging, and cancer^39–50^. Interestingly, Stéphane Pelletier showed that GPCR agonists stimulate tyrosine phosphorylation of STAT3 proteins in a Rac-dependent manner^51^. Shangwen Liu et al found the functions of Notch1 are responsible for mediating PTEN/PI2K/AKT pathway in renal cancers^52^. Necip Pirinççi et al found the serum CAT activity was significantly decreased in patients with renal cancers by comparing the controls^53^. As a tumor biomarker, the upregulation of cyclin D1 has the role of predicting the favorable prognosis in patients with renal cancers^54^. Jun Li et al found HSPA8 is a potential prognostic factor to predict the survival of patients with AML^55^. Yun Ye et al found DLG4 is a potential prostate cancer marker by analyzing the mRNA-miRNA-lncRNA database^56^. Shangqing Song et al found the urinary exosome miR-30c-5p can inhibit the progression of renal cancers by targeting HSPA5^57^.

In conclusion, this study found the significant genes and biological processes between the 2D and 3D culture methods in renal cancer cells. The “sulfur compound metabolic process” and “regulation of leukocyte mediated immunity” are the top affected signalings between the different culture methods. Our study may provide novel insights for the treatment of renal cancers in vitro.

## Author Contributions

Mengyao Wang, Hongmei Guo: Methodology and Writing. Hanming Gu, Mason Zhang: Conceptualization, Writing-Reviewing and Editing.

